# TMS with fast and accurate electronic control: measuring the orientation sensitivity of corticomotor pathways

**DOI:** 10.1101/2021.08.20.457096

**Authors:** Victor Hugo Souza, Jaakko O. Nieminen, Sergei Tugin, Lari M. Koponen, Oswaldo Baffa, Risto J. Ilmoniemi

## Abstract

**Background:** Transcranial magnetic stimulation (TMS) coils allow only a slow, mechanical adjustment of the stimulating electric field (E-field) orientation in the cerebral tissue. Fast E-field control is needed to synchronize the stimulation with the ongoing brain activity. Also, empirical models that fully describe the relationship between evoked responses and the stimulus orientation and intensity are still missing.

**Objective:** We aimed to (1) develop a TMS transducer for manipulating the E-field orientation electronically with high accuracy at the neuronally meaningful millisecond-level time scale and (2) devise and validate a physiologically based model describing the orientation selectivity of neuronal excitability.

**Methods:** We designed and manufactured a two-coil TMS transducer. The coil windings were computed with a minimum-energy optimization procedure, and the transducer was controlled with our custom-made electronics. The electronic E-field control was verified with a TMS characterizer. The motor evoked potential amplitude and latency of a hand muscle were mapped in 3° steps of the stimulus orientation in 16 healthy subjects for three stimulation intensities. We fitted a logistic model to the motor response amplitude.

**Results:** The two-coil TMS transducer allows one to manipulate the pulse orientation accurately without manual coil movement. The motor response amplitude followed a logistic function of the stimulus orientation; this dependency was strongly affected by the stimulus intensity.

**Conclusion:** The developed electronic control of the E-field orientation allows exploring new stimulation paradigms and probing neuronal mechanisms. The presented model helps to disentangle the neuronal mechanisms of brain function and guide future non-invasive stimulation protocols.

## Introduction

Transcranial magnetic stimulation (TMS) has been used extensively to study human brain function and treat many neurological diseases [1]. In TMS, the intensity and orientation of the induced electric field (E-field) are critical factors to excite specific neuronal populations [2–5] and to activate distinct neurotransmitters [6,7]. Conventionally, the E-field orientation for an optimal response is adjusted manually—a slow process required in many TMS applications. Manual TMS control does not allow manipulating the E-field orientation at the millisecond-level time scale of cortical signaling, which is needed for probing neuronal excitatory and inhibitory processes with different stimulation orientations [8,9]. Even when assisted by neuronavigation, manual scanning of the optimal coil orientation has a relatively high error associated with the coil placement [10,11], resulting in reduced cortical specificity. Electronic control of the E-field orientation would enable automated experimental procedures and new stimulation protocols for studying neuronal processes with TMS.

Electronic control of the stimulus location and orientation on the cortex can be achieved by independently generated TMS pulses with coils placed on the scalp in a multi-coil TMS (mTMS) configuration [12–14]. The mTMS concept was introduced by Ruohonen et al. [13,15]; however, the originally proposed coil arrays would require many coils to fine control the peak induced E-field [15] and be associated with excessive power electronics requirements. To overcome the limitations, we developed a minimum-energy algorithm to design optimal TMS coils [16] and developed an mTMS transducer comprising two overlapping coils (a figure-of-eight and an oval coil) capable of electronically shifting the stimulation locus along a line segment in the cortex [12]. We have also developed an mTMS system with a large general-purpose 5-coil transducer [14]. Previous studies have also combined two figure-of-eight coils to produce a rotating E-field in a single TMS pulse capable of reducing the orientation sensitivity of neuronal excitation [17,18]. In this study, we developed a compact 2-coil transducer tailored for controlling the orientation of the peak induced E-field to allow high-resolution mappings of the neuronal sensitivity to the TMS pulse orientation.

Changing the stimulus orientation for the excitation of a neuronal population, for instance, pyramidal neurons with anisotropic dendritic arborization, is perceived fundamentally as a variation in the effective stimulation intensity [19]. The induced E-field parallel to the longitudinal axis of pyramidal neurons, i.e., along the cortical columns, leads to maximal excitation of the targeted neurons, as predicted by the so-called cortical column cosine model [20]. On a macroscopic scale, this model implies that the stimulation effect is proportional to the cosine of the angle between the E-field and the normal of the cortical surface. The cosine model has been largely employed in simulations to estimate the effect of TMS in the brain [21,22]. On the other hand, recent computational and experimental studies have suggested that TMS excites neurons mostly at axon terminals and that all components of the E-field, i.e., not only its normal component, have a substantial contribution to the neuronal depolarization [23–25]. Although previous results have advanced our understanding of TMS-triggered cortical activation, empirical models that fully describe the relationship between the evoked response and the stimulus orientation and intensity are still missing.

We aimed to develop an mTMS transducer that allows an experimenter to rotate the peak E-field electronically, i.e., without manual coil movement. With the help of the new transducer, we aimed to measure how the motor evoked potentials (MEPs) depend on the stimulus orientation and intensity, emphasizing the importance of fine E-field adjustments to unveil the neuronal effects of TMS.

## Material and Methods

### mTMS transducer

We designed and built a transducer comprising a pair of orthogonally oriented figure-of-eight coils using our minimum-energy optimization method [12,16]. This method provides energy-efficient coil windings, reducing the power electronics requirements and heat dissipation in the coils. First, we modeled a commercial figure-of-eight coil (70-mm Double Coil; The Magstim Co Ltd, UK; **Fig. 1A** and **B**) [26] placed 15 mm above a spherical cortex model (70-mm radius; 2562 points on the surface). Then, we computed a set of reference E-field distributions by rotating the commercial coil from 0° to 180° in steps of 10° around the normal of the cortex and the coil bottom. The E-field distributions were computed using an analytical closed-form solution and reciprocity [12,27,28]. Next, for each reference E-field, we calculated the minimum-energy surface current density in an octagonal plane section (15-cm outer diameter; 1089-vertex triangular mesh) by minimizing the magnetic energy of the surface current density to generate an E-field similar (the same peak amplitude and the same focality) to the reference E-field distribution [12,16]. The focality was quantified by the extent of the cortical region where the E-field magnitude exceeded 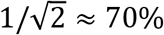 of its maximum. The octagonal coil geometry provides identical boundaries for windings spanning in all directions of any of two perpendicular lines of symmetry, as opposed to, e.g., a hexagonal geometry, which has different boundaries along two perpendicular lines of symmetry. This property helps in preventing different truncations of the winding paths for the bottom and top coils.

**Fig. 1.**
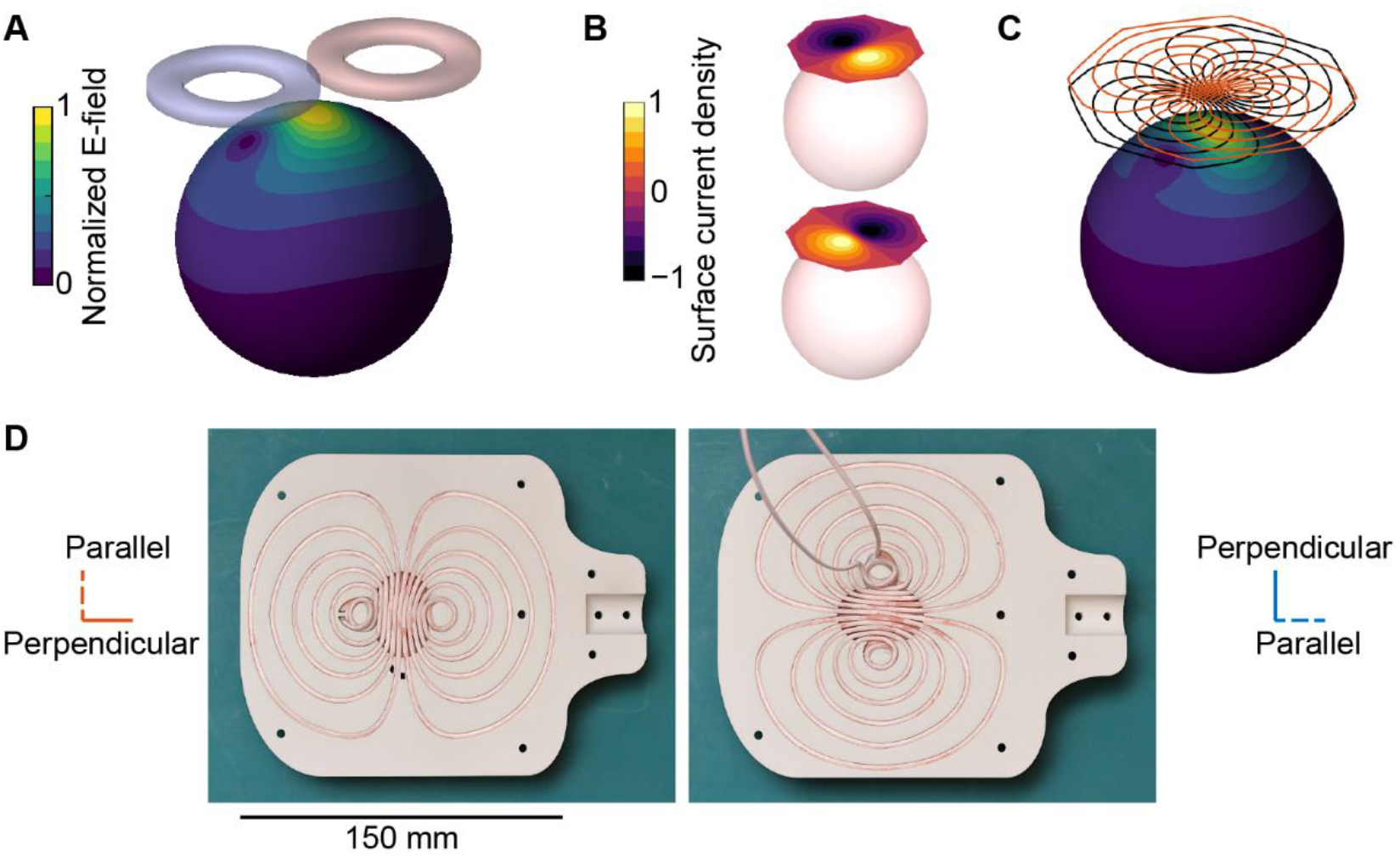
mTMS transducer. A) Example of a target E-field induced on a spherical surface by a model of a commercial figure-of-eight coil. B) Optimized minimum-energy surface current density distributions for the bottom and top coils. C) Surface current distributions of (B) discretized in 12 turns of wire. The induced E-field has a focality similar to that of the figure-of-eight coil in (A) but can be rotated by adjusting the currents in the top (orange) and bottom (black) coils. D) mTMS transducer with litz wire wound in the bottom and top 3D-printed formers. The two coils have identical windings but are assembled to induce E-fields in perpendicular orientations.

The resulting set of surface current densities was decomposed with the singular value decomposition, with each component representing a single coil. The winding paths for the two selected coils were optimized separately using planes placed 15 and 20 mm above the cortical surface, respectively. Finally, we computed the contour lines of these two surface current distributions to obtain the paths for the coil windings (**Fig. 1C**).

The coil formers were designed in SolidWorks 2016 (Dassault Systèmes SA, France) and printed by selective laser sintering of 30% glass-filled polyamide (Maker 3D, Finland). Glass-filled polyamide (tensile strength: 38 MPa; dielectric strength: 15 kV/mm) is resistant to the pressure of about 10 MPa from the Lorentz forces during the TMS pulse (derived from [29]). We wound in series two layers of 12 turns of copper litz wire (1.6 mm thick and 2.4 mm wide; Rudolf Pack GmbH & Co. KG, Germany) to each former (**Fig. 1D**). The transducer assembly was potted with epoxy for increased strength and electrical insulation, and each coil was soldered to a cable provided by Nexstim Plc (Finland).

The electronically rotated E-field distribution was measured at 1000 points in a hemisphere (70-mm radius) for stimulus orientations of 0°, 45°, and 90° with our TMS characterizer [30]. In this study, the term stimulus orientation refers to the direction of the peak induced E-field in a spherical head model. With the TMS characterizer, we measured the profiles parallel and perpendicular to the peak E-field below the coil center and computed the focality as the width of the region where the E-field exceeded 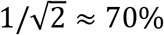 of its maximum [16,30]. The mTMS pulses were monophasic and delivered by our custom-made electronics [12,31], with the waveform divided into three parts lasting for 60.0 μs (rising), 30.0 μs (holding), and 43.2 μs (falling) [31].

The current pulse waveform was measured for each coil with a Rogowski probe (CWT 60B; Power Electronic Measurements Ltd, UK) connected to an oscilloscope (InfiniiVision MSOX3034T; Keysight, USA). The self-inductance and resistance of each coil were determined by connecting the coil in series with a half-bridge circuit with a 99.3-Ω resistor and measuring the phase difference between the voltage across the coil and the half-bridge. The circuit was powered with a sinewave voltage at frequencies of 1, 2, 3, 5, 10, 20, and 50 kHz (AFG1062, Tektronix, USA) and voltages were measured with an oscilloscope (Keysight).

### Participants

Sixteen healthy subjects (mean age: 29 years, range 22–41; five women) participated in the study. All participants gave written informed consent before their participation. The study was performed in accordance with the Declaration of Helsinki and approved by the Coordinating Ethics Committee of the Hospital District of Helsinki and Uusimaa.

### Experimental procedure

Subjects sitting in a reclining chair were instructed to stay relaxed during the TMS sessions. Electromyography (EMG) was recorded from the right abductor pollicis brevis (APB) using surface electrodes in a belly—tendon montage (Neuroline 720, Model 72001-K/12; Ambu, Denmark) with a Nexstim eXimia EMG device (500-Hz lowpass filtering, 3000-Hz sampling frequency; Nexstim Plc). The mTMS transducer was placed over the left motor cortex, guided by the individual cortical anatomy using a neuronavigation system (NBS 3.2, Nexstim Plc). Anatomical T1 magnetic resonance imaging was performed before the mTMS experiments (voxel dimensions less than or equal to 1 mm). The mTMS pulse waveforms were the same as those used for the transducer calibration.

First, the APB hotspot was identified as the cortical site beneath the transducer center resulting in MEPs with the maximum peak-to-peak amplitude for single-pulse TMS. The hotspot was obtained with the peak E-field induced by the bottom coil being approximately perpendicular to the central sulcus and with its first phase inducing an E-field from posterolateral to anteromedial direction. After defining the hotspot, the transducer was manually rotated to obtain the highest MEP amplitudes with a fixed suprathreshold intensity. Then, the resting motor threshold (MT) was estimated as the minimum stimulation intensity eliciting at least 10 out of 20 MEPs with at least 50 μV of peak-to-peak amplitude [1]. The MT was estimated for stimulation at 0° (MT0°; posterolateral to anteromedial) and 90° (MT90°; posterolateral to anteromedial) E-field orientations (see **Fig. 4A** for a schematic representation of the E-field orientations relative to the subject’s head).

To measure the effect of the E-field orientation on the MEP amplitude and onset latency, we applied five single pulses to the APB hotspot of 11 subjects with 110% of the MT_0°_ at each orientation in 3° steps, i.e., 120 orientations. The interval between consecutive pulses was pseudo-randomized from 4 to 6 s sampled from a uniform distribution. To investigate the effect of the stimulation strength on the orientation dependency, we applied a similar experimental protocol on five other subjects. MEPs were collected from three single pulses at the APB hotspot at each orientation in 3° steps (i.e., 120 orientations) for each of the three stimulation intensities 110% and 140% of MT_0°_, and 120% of MT_90°_. For two out of these five subjects, we measured the MEPs for an additional intensity of 120% of MT_0°_. The interval between consecutive pulses was pseudo-randomized from 2.4 to 2.7 s sampled from a uniform distribution. All pulses were divided into sequences lasting approximately 6 min each, followed by short breaks of about 5 min. The order of the orientation and intensity of the pulses was also pseudo-randomized.

### Data processing

Using custom-made scripts written in MATLAB R2017a (MathWorks Inc, USA), MEPs were extracted from the continuous EMG recordings, and trials showing muscle pre-activation or movement artifacts greater than ±15 μV within 1000 ms before the TMS pulse were discarded. For each trial, we computed the MEP peak-to-peak amplitude in a time window 15–60 ms after the TMS pulse and manually annotated the MEP latency. Given the inherent uncertainty in assigning the latency to small MEPs, the latencies from trials with a peak-to-peak amplitude below 50 μV were not included in the analysis (0.9% of the trials were discarded and 51% of the MEP latencies were not annotated). As expected, the relatively high amount of non-annotated latencies was due to the small MEPs around the ±45 and ±135° orientations.

To align the data across subjects, we smoothed each subject’s orientation–amplitude curve with a moving-average filter with a window size of 7 (i.e., considering data for pulses within 18°) and fit a two-peak composite Gaussian to extract the peak orientation. The angle corresponding to the maximum amplitude was subtracted from the original angles, ensuring that all subjects had the maximum amplitude at 0°. The average absolute correction across subjects was 8.5° (standard deviation: 6.5°). The correction obtained from the amplitude data was applied to the latency data, maintaining the correspondence between both measures.

### Modeling the orientation selectivity of neuronal excitation

First, we visually assessed how the cosine [20] and the Gaussian equation [32] fit the measured MEP amplitudes versus stimulus orientations. These models failed to capture the essential characteristics of the entire peak, as shown in **Fig. 2**. Note that the cosine did not fit well to the exponential increase in the MEP amplitude between 45° and 90°. The Gaussian model did not match the inflection around ±45° and the plateau around the peak due to the sharp increase in the MEP amplitude data from the baseline, followed by a smoother approach to the peak. The logistic (sigmoid) equation, on the other hand, describes well the relationship between the stimulus strength (input) and the neuronal output [33], making it a suitable candidate to parametrize the orientation dependency of the corticomotor response. However, the logistic function changes monotonically and, in principle, is unsuitable to model peak-shaped curves. To overcome this issue, we divided each observed peak, centered at 0° or 180°, in two quadrants resulting in segments with monotonically increasing (−90° to 0°; 90° to 180°) or decreasing (0° to 90°; 180° to −90°) MEP amplitudes. We modeled the log-transformed MEP amplitude [33,34] versus the stimulus orientation with the generalized logistic equation below, for each quadrant (−180° to −90°; −90° to 0°; 0° to 90°; 90° to 180°) and subject.

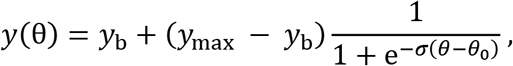

where 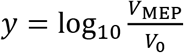, *V_MEP_* is the MEP amplitude, *V*_0_ = 1 μV a reference MEP amplitude (to obtain a unitless argument for the logarithmic function), *y*_b_ the baseline MEP amplitude, *y*_max_ the maximum MEP amplitude, *θ* the stimulus orientation, and *θ*_0_ and *σ* the center and slope of the logistic equation, respectively. Considering data continuity on the closed circle (−180° to 180°), we extended each quadrant by 15° (5 samples) on both sides to minimize the edge effects when computing the fit. The extra samples were removed after the model parameters were calculated. For the statistical analysis, we classified the maximum (*y*_max_) and baseline (*y*_b_) amplitudes from the −90° to 0° and 0° to 90° quadrants as belonging to the 0° peak. Similarly, *y*_max_ and *y*_b_ from the −180° to −90° and 90° to 180° quadrants were classified as the 180° peak (results reported in **Fig. 5A–D**).

**Fig. 2.**
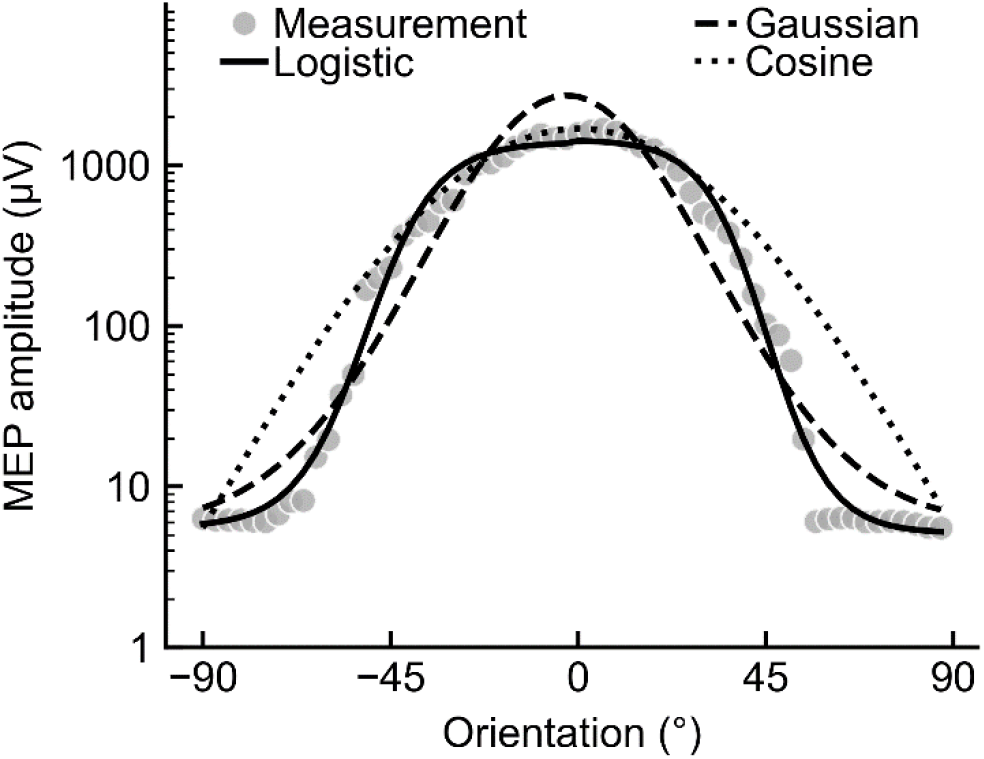
Modeling the MEP amplitude versus stimulus orientation. Comparison between the logistic equation (solid line), cosine (dotted line), and Gaussian function (dashed line). The gray markers are the median MEP amplitudes across three trials recorded from one subject (S15) with stimulation at 110% MT_0°_.

The logistic model parameters can provide information on the structural–functional relationship of the primary motor cortex. Given a set of pre-defined physical constraints, such as the pulse shape, coil-to-cortex distance, and intensity, *y*_max_ represents the neuronal capacity as the maximal response that can be obtained from the stimulated population and *y*_b_ the corresponding minimum neuronal response. σ represents the orientation sensitivity of the neuronal population; high values indicate a neuronal population preferentially aligned in a narrow range of orientations resulting in a sharp change between the baseline and maximum amplitudes. Lastly, *θ_0_* characterizes the intrinsic properties of the neuronal population combining *y*_max_, *y*_b_, and *σ*.

For the MEP latency, only subjects showing clear responses across the entire range of orientations for each intensity were included in this analysis (7 subjects in 110% MT_0°_, and five subjects in 140% MT_0°_ and 120% MT_90°_). Then, we fit a 2^nd^-order real trigonometric polynomial equation on the latency data set of each stimulation intensity and subject, accounting for the periodicity and visible asymmetry of the data across the closed circle:

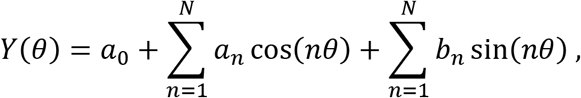

where *Y*(0) is the measured latency as a function of the stimulation angle *θ* in radians, *N* = 2 the degree of the trigonometric polynomial, and *a_n_* and *b_n_* (0 ≤ *n* ≤ *N* and *a_N_* ≠ 0 or *b_N_* ≠ 0) the regression coefficients. From the fitted data, we extracted the MEP latency at −90°, 0°, 90°, and 180° for the statistical analysis (results reported in **Fig. 5E**).

The regression residuals were visually inspected for deviations from normality. For modeling, we used the least-squares solver implemented in the *lmfit* 1.0 package, and for data pre-processing and visualization, we used custom-made scripts written in Python 3.7 (Python Software Foundation, USA).

### Statistical analysis

We applied a linear mixed-effects model to assess the effect of stimulus intensity and peak orientation on the MEP amplitude model parameters and latency. The intensity and peak orientation were modeled as fixed effects, while subjects were modeled as a random effect using restricted maximum likelihood estimation. The *p*-values of the fixed effects were derived with Satterthwaite approximations in a Type III Analysis of Variance table. Post-hoc multiple comparisons were performed using the estimated marginal means with *p*-value correction for the false discovery rate. Further description of the statistical analysis is provided in the *Supplementary Material*. The threshold for statistical significance was set at *p* = 0.05.

## Results

### mTMS transducer to rotate the E-field electronically

The measured E-field distributions with the stimulus orientation electronically rotated to 0°, 45°, and 90° are illustrated in **Fig. 3A**. The stimulation intensity in every orientation reached up to 129 V/m (average E-field during the first part of the pulse waveform) measured with our probe at 70 mm from the center of an 85-mm-radius spherical head model. The E-field profiles for the bottom and top coils are illustrated in **Fig. 3B**. In the direction perpendicular to the peak E-field, the focalities for the bottom and top coils were 25.6 and 26.5 mm, respectively; in the parallel direction, they were 44.8 and 46.4 mm, respectively. The current required by each coil follows the cosine and sine function of the desired E-field orientation and norm, as expected, given that the coils are orthogonal. **Fig. 3C** shows the monophasic current and induced E-field waveforms for both coils. The bottom and top coils required 54 A/μs (868 V) and 70 A/μs (1128 V), respectively, to induce a 100 V/m E-field measured by our probe [30]. The average deviations from the theoretical values in the E-field norm for stimulation at 25 V/m and orientation from 0° to 180° were 0.1 V/m and 1.3°, respectively. The mean (min; max) measured self-inductance and resistance across the tested sinewave frequencies were, respectively, 12.9 (12.0; 14.8) μH and 87.1 (72.8; 99.6) mΩ for the bottom coil, and 13.1 (12.2; 15.1) μH and 95.3 (72.1; 100.6) mΩ for the top coil. Reasons for the dependence of these values on frequency are the skin effect and the proximity effect.

**Fig. 3.**
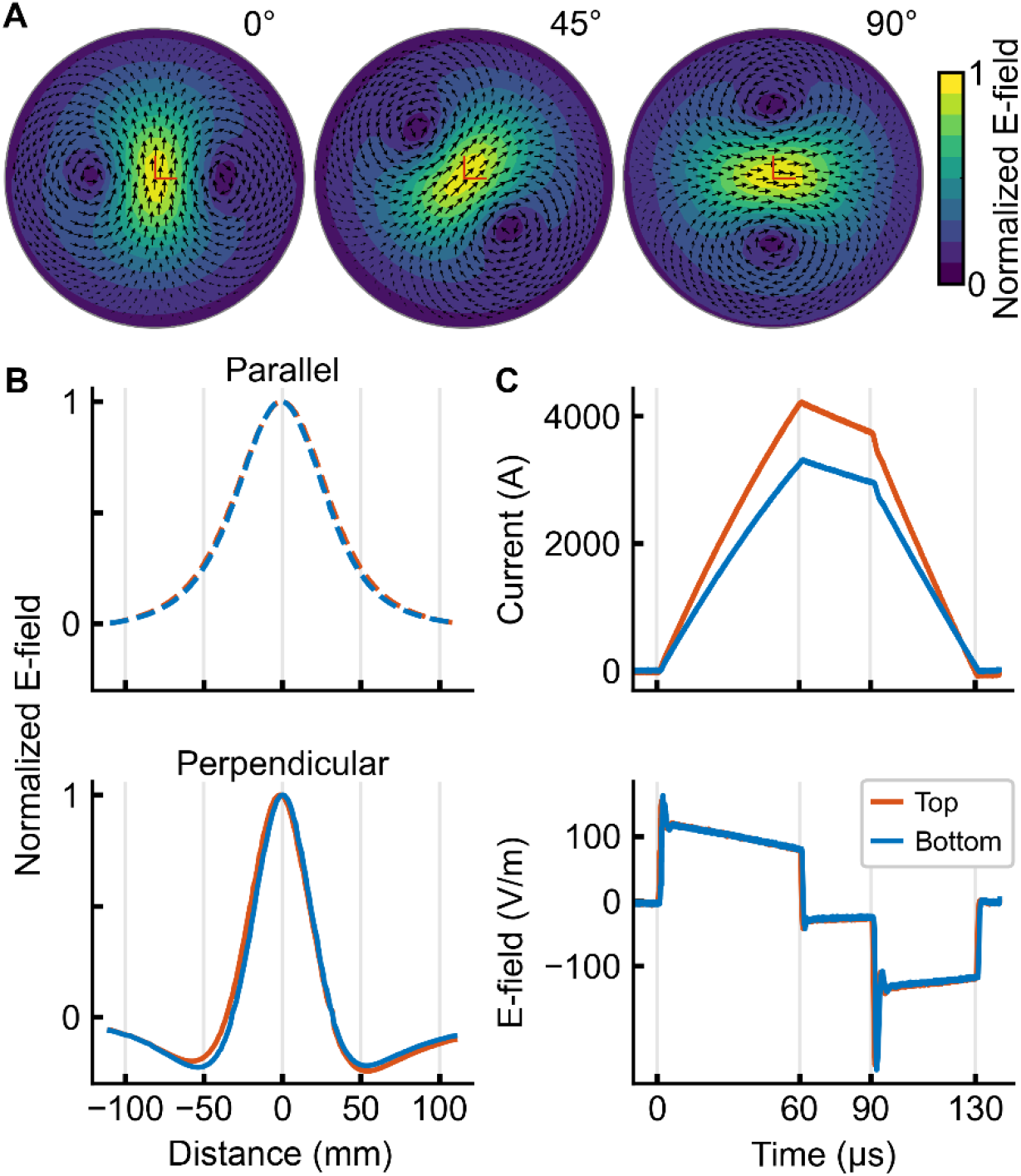
mTMS E-field, focality, and current. A) Measured E-field distribution induced on a spherical surface with a 70-mm radius for a TMS pulse at 0°, 45°, and 90°. B) The profile along the direction parallel and perpendicular to the peak induced E-field, respectively. The solid and dashed lines refer to the bottom and top coils, respectively. C) Monophasic current waveforms (upper panel) from the bottom and top coils and the corresponding induced E-field waveforms (lower panel) with a 100-V/m (on average) E-field during the rising part of the current. The top coil requires a higher current to induce the same peak E-field intensity due to the increased distance from the cortex. The waveforms have been lowpass filtered at 1 MHz.

**Fig. 4.**
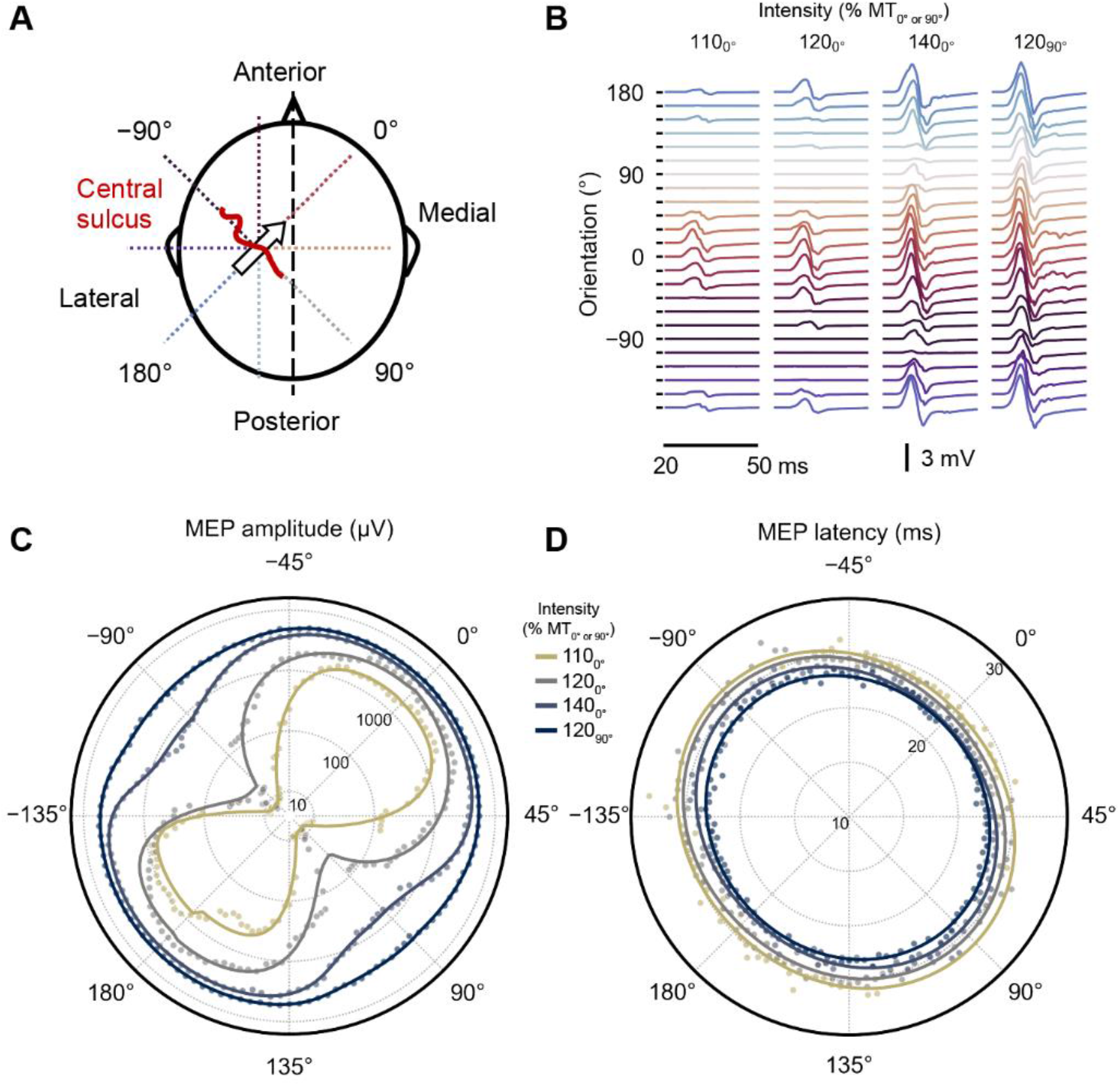
MEP amplitude and latency as functions of the stimulus orientation and intensity. A) A schematic representation of the stimulus orientations relative to the subject’s head. The 0° orientation refers to the first phase of the induced E-field pointing to the anteromedial orientation, approximately perpendicular to the central sulcus. B) The MEP epochs (median across three trials), from 20 to 50 ms after the TMS pulse, at each stimulus orientation and intensity from S16. The colors encode the stimulus orientation depicted in (A). The logistic and trigonometric polynomial fits (solid lines) of the C) MEP amplitude and D) latency, respectively, for each intensity as a function of the stimulus orientation for S16. The radial MEP amplitude axis is logarithmic. The dots represent the median MEP amplitude and latency across the repeated trials for each stimulus orientation and intensity. The colors indicate the stimulation intensity in % of the resting motor threshold at 0° (MT_0°_) and 90° (MT_90°_).

**Fig. 5.**
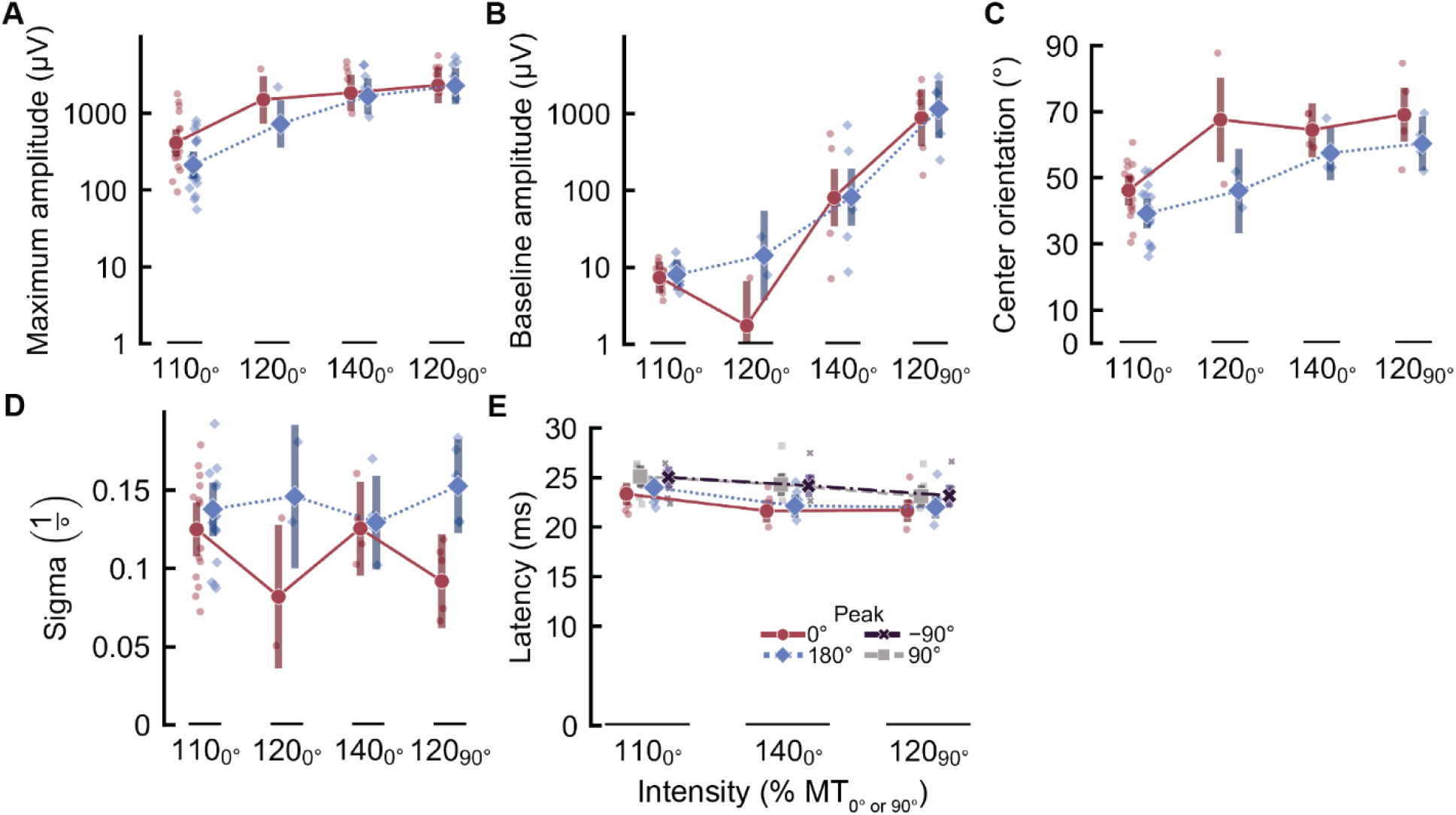
MEP amplitude and latency parameters. Linear mixed model analysis of the logistic equation parameters A) maximum amplitude (*y*_max_), B) baseline amplitude (*y*_b_), C) center orientation (*θ*_0_), and D) slope sigma (*σ*) across subjects. The *y*_max_ and *y*_b_ of the 0° and 180° peaks were extracted from the (−90° to 0°; 0° to 90°) and (−180° to −90°; 90° to 180°) quadrants, respectively (see the section *Modeling the orientation selectivity of neuronal excitation*). The error bars represent the 95% confidence interval of the estimated marginal means. The difference in error bar sizes is due to the distinct number of subjects in each intensity (16 subjects in 110% MT_0°_, two subjects in 120% MT0°, and five subjects in 140% MT_0°_ and 120% MT_90°_). E) Linear mixed-effects model analysis of the MEP latency extracted from the four peak orientations (−90°, 0°, 90°, and 180°) in the trigonometric polynomial fit. We removed the MEP latency from 120% MT_0°_ because only subjects showing latencies across the entire range of orientations for each intensity were included in this analysis (7 subjects in 110% MT_0°_, and five subjects in 140% MT_0°_ and 120% MT_90°_).

### Effect of E-field orientation and intensity on motor response

The orientation dependency of the MEP amplitude was greatly affected by the increase in the stimulation intensity. We observed the same effect in all five subjects, illustrated with data from subject 16 (S16) in **Fig. 4B, C**, and **D**. The traces in **Fig. 4B** show two clusters of motor responses around 0° and 180° with increasing response amplitude for the stimulus intensities 110%, 120%, and 140% of the MT_0°_. At 120% of the MT_90°_, all orientations seem to generate MEPs with similar amplitudes. Data for all the subjects are provided in *Supplementary Figures 1* and *2*.

The logistic equation captured well the changes in the median MEP amplitude as a function of the orientation and intensity as shown in **Fig. 4C** for S16 (plots for all subjects are provided in *Supplementary Figure 1*). The linear mixed-effects model and multiple comparison results are all provided in *Supplementary Tables 1–15*. At 110% of the MT_0°_, we observed a maximum MEP amplitude about two times greater at 0° than at 180°(*df* = 33.0; *t*_ratio_ = 4.78;*p* < 0.001; **Fig. 5A**), with the peak E-field approximately perpendicular to the central sulcus but pointing in opposite directions. Such difference was not evident between these orientations at the two highest intensities (140% MT_0°_: *df* = 33.0; *t*_ratio_ = 1.11; *p* = 0.73; 120% MT_90°_: *df* = 33.0; *t*_ratio_ = 1.02; *p* = 0.94; **Fig. 5A**). The maximum MEP amplitude reached the saturation value at a lower intensity (120% of the MT_0°_) than the baseline amplitude, which only increased with the tested stimulation intensities (**Fig. 5A** and **B**). This was expected due to the 28% higher MT at 90° compared with 0° orientation (one-tailed paired *t*-test, *p* < 0.001). Thus, the difference between the maximum amplitude, at 0° or 180°, and the baseline amplitude, at ±90°, greatly reduced with the increase in the stimulation intensity. The highest stimulation intensities also led to similar MEP amplitudes across all orientations, evidenced by the change from an eight-shaped curve at 110% MT_0°_ to a nearly circular one, with almost constant amplitude at 120% of MT_90°_ (**Fig. 4C**). The center orientation increased linearly with the intensity (**Fig. 5C**), corresponding to the broader peaks in the orientation dependency curve, as illustrated for S16 (**Figs. 4C**). In turn, the logistic equation slope (sigma) did not show a clear variation with the stimulation intensity apart from a small difference between the 0° and 180° peaks at 120% of MT_90°_ (*df* = 34.0; *t*_ratio_ = −3.27;*p*< 0.039; **Fig. 5D**).

For all stimulation intensities, MEP latencies at 0° and 180° were about 2 ms shorter than those at 90° and 270° (**Fig. 5E**). Moreover, the lowest stimulation intensity (110% MT0°) evoked MEPs with about 2 ms longer latencies than stimulation at higher intensities (140% MT_0°_ and 120% MT_90°_) at 0° or 180° orientation. The MEP latency was 0.7 ms longer at 180° than at 0° with the stimulus intensity at 110% of MT_0°_ (*df* = 47.0; *t*_ratio_ = −2.85;*p* < 0.008). In turn, at 140% of MT_0°_ and 120% of MT_90°_, we did not observe significantly different MEP latencies between stimuli at 0° and 180°.

## Discussion

We developed a 2-coil TMS transducer that can control the E-field orientation electronically at millisecond-scale intervals without a need to rotate the transducer. The millisecond-interval targeting can be achieved by adjusting the current waveforms without recharging the capacitors [8,35]. The electronic control of the stimulus orientation enables an accurate adjustment of the TMS pulse orientation by automated algorithms [36,37] in closed-loop paradigms triggered by neurophysiological recordings [38,39]. The automated protocols may increase the efficacy of TMS by maximizing the cortical excitation through the optimal alignment between the induced E-field and the neuronal populations. The fast control of stimulus orientation also allows advanced studies on intracortical inhibition and facilitation mechanisms [6,7,9]; for instance, in paired-pulse protocols, the system can change the E-field orientation within a millisecond interval without the need for the mechanical rotation of the transducer. Moreover, we demonstrated that the simple logistic equation describes the motor response dependency on the stimulus orientation and intensity better than just the cosine of the peak E-field orientation, fostering the development of more accurate and physiologically based computational models of the neuronal effects of TMS. This model can also be used as *a priori* information when automatically searching for the optimal E-field orientation, making the procedure possible with fewer pulses than without the model.

### mTMS transducer

The 2-coil transducer has a size similar to that of commercial figure-of-eight TMS coils and only half the size (150 mm) of our other mTMS transducers: the 2-coil transducer for shifting the stimulation locus within a line segment [12] and the 5-coil mTMS transducer that can shift and rotate the peak E-field within a cortical region [14]. The recently developed 5-coil mTMS transducer comprises a similar combination of two perpendicular figure-of-eight coils to rotate the E-field orientation electronically; however, the large size (300 mm × 300 mm) prevents its use in cortical areas with restricted space. The compact design of the present 2-coil transducer offers more comfortable positioning over the scalp and better handling, required for many applications outside the motor cortex, such as in the visual cortices [40], cerebellum [41], and dorsolateral prefrontal areas [42].

The required energy depends on the stimulation orientation. For example, to induce a 100-V/m E-field at a depth of 15 mm from the surface of an 85-mm radius sphere, the bottom coil requires 45 J, which is 41% lower than the energy required by the top coil (76 J) and 12%higher than the energy needed with an optimized 300-mm-wide figure-of-eight coil (40 J) [43]. In other terms, the transducer’s bottom coil requires 23% lower current (and voltage) than the top coil to generate an equally strong E-field at a depth of 15 mm from the surface of an 85-mm radius sphere. This difference is due to the top coil’s 5-mm extra distance from the cortical surface, which leads to a weaker coil–cortex coupling [30,43]. The increased distance also leads to 3% (1–2 mm) poorer focality in perpendicular and parallel directions for the top compared to the bottom coil, a negligible difference for standard TMS applications. If desired, this difference can be eliminated in future models by designing the bottom coil so that its focality matches the other coil’s pattern [44]. The possibility to use 3D-printed formers might ease the manufacturing of transducers with more complicated winding patterns, such as those reported by Deng et al. [45] and mTMS transducers with more coils [12,14].

### Orientation selectivity of the TMS effect

Our measurements revealed that, for relatively weak stimuli, the MEP amplitude abruptly decreases when moving away from the optimal orientation, with a stronger dependency than just the first-degree cosine of the angle. We observed a single, smooth amplitude peak at the optimal orientation for all 16 participants, in contrast to the bimodal response curves at 0° reported for most participants by Kallioniemi et al. [32]. The bimodal response might have been caused by inaccuracies inherent to the manual coil adjustments, which are eliminated by the electronic rotation of the E-field with the 2-coil mTMS.

The increase in stimulation intensity severely reduced the orientation selectivity of the TMS effect. One explanation is that distinct neuronal populations with different excitation thresholds are recruited depending on the stimulus orientation [46,47]. For instance, if we assume that on the hand knob area of the primary motor cortex, a neuronal population P_1_ is optimally aligned with the 0° stimulus while a population P_2_, slightly further away from the hotspot, is aligned at 90°, illustrated in **Fig. 6**; a low-intensity stimulation delivered at 0° would selectively excite population P1 while the same intensity but at a 90° orientation would not reach the excitation threshold of population P_2_. An increase in the intensity would rapidly lead to the maximal MEP amplitude of population P_1_ (0° stimulus and saturation level in the input-output curve) while only slightly above the excitation threshold of population P_2_ (90° stimulus). Further increasing the stimulation intensity would excite P_2_ (90° stimulus) more without further effect on P_1_ (0° stimulus), which is already at maximum capacity.

**Fig. 6.**
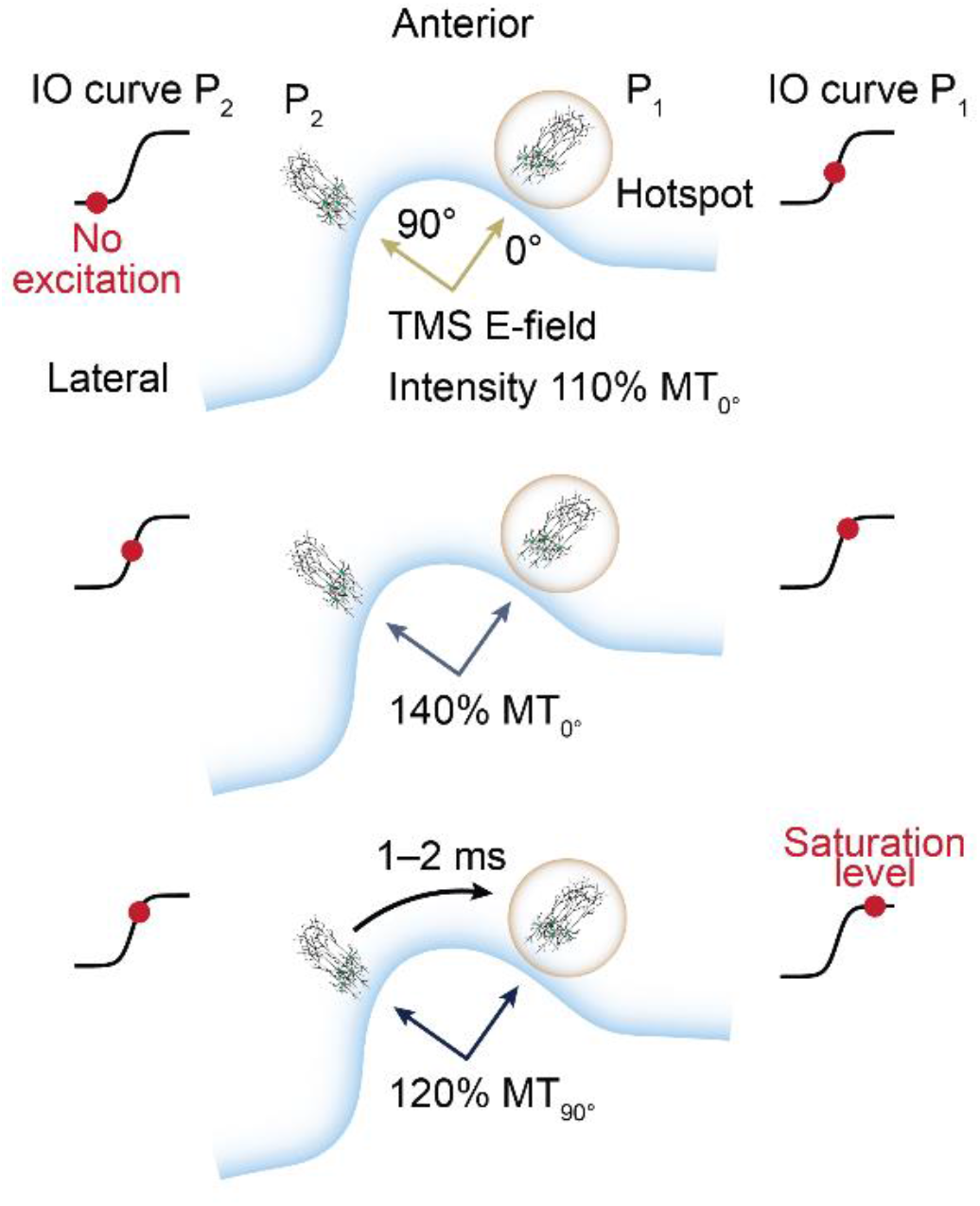
Hypothetical mechanisms of cortical excitation due to distinct TMS-induced E-field orientations and intensities. A putative model for the effect of stimulation intensity in the excitation of neuronal populations P_1_ (hotspot) and P_2_ (vicinity) by the two perpendicular E-field orientations (0° and 90°). The level of neural output depends on each population’s input–output (IO) curve and the relative alignment with the peak E-field orientation.

Our findings support this activation model from multiple perspectives. First, the MEP amplitude induced with stimuli at ±90° rapidly increased to a similar level to those obtained at 0°, evidenced by the steeper increase in the baseline amplitude compared to the maximum amplitude (**Fig. 5A** and **B**). Second, the center orientation increased linearly with the intensity, indicating a reduction in the orientation selectivity (**Fig. 5C**). Lastly, the MEP latency was about 1.5–2.6 ms shorter at 0° compared with the ±90° orientations (**Fig. 5E**). Such delay would justify the excitation of the neuronal population P_2_ in the vicinity of the cortical hotspot, projecting to the hotspot through one synaptic connection. In fact, we showed in a recent study that neuronal populations near the cortical target site seem to affect the generated neural drive [35].

Surprisingly, the slope (σ) of the orientation dependency curve did not show a clear dependence on the stimulation intensity. This is probably due to the proportional, simultaneous change in the center orientation (*θ*_0_) and the baseline amplitude (*y*_b_), combining the characteristics of all the excited neuronal populations in all directions in the orientation dependency slope. The input–output curve slope has been shown to depend on the TMS pulse shape and waveform, revealing properties of the membrane depolarization [33]. However, changing the stimulus orientation may excite neuronal populations in different cortical sites, as discussed above, hindering the discrimination of cellular properties from a specific neuronal ensemble through the orientation dependency curve slope.

The MEPs are modulated by both cortical and subcortical (cerebellar and spinal) effects [4,48]. It appears to be an oversimplification to assume that the neuronal or muscle activation is proportional only to the cosine between the induced E-field and the pyramidal neurons’ somatodendritic axis, as previously reported [20,21]. The updated view is supported by recent multi-scale realistic simulations, in which all E-field components, rather than only the normal component, seem to contribute significantly to the neuronal excitation due to the widespread axonal ramification in different orientations [24]. Furthermore, the excitatory and inhibitory interneurons contributing to the cortical excitation and its control exhibit specific alignments relative to the cortical columns [49,50]. For instance, excitatory neurons mainly project from layers 2 and 3 to pyramidal neurons in layer 5, while inhibitory interneurons exhibit a stellate arborization with mainly horizontal projections within layers 2 and 3 [51]. In this case, inhibitory mechanisms have a lower excitation threshold and are less affected by the E-field orientation [6], possibly explaining the stronger decay in the muscle response in suboptimal directions. Thus, our models provide evidence on the structure–function relationship of neuronal populations from the primary motor cortex, encompassing all cortical mechanisms engaged in the MEP generation.

We found that MEP latencies are shortest for the 0° stimulus, i.e., when the E-field is perpendicular to the cortex, and somewhat longer for the ±90° stimulus for all stimulation intensities. Changes in latency presumably reflect differences in the generation of cortical direct and indirect descending volleys. In contrast to our observations, previous studies showed that monophasic TMS pulses delivered perpendicular to the central sulcus elicited stronger indirect waves (I-waves), i.e., long-latency MEPs. In turn, pulses along the central sulcus have been shown to produce stronger direct (D-) waves resulting in short-latency MEPs [47]. One explanation is that the optimal orientation (around 0°) evoked earlier descending volleys I_1_–I_3_ [52], while 90° pulses preferentially recruited later waves, e.g., I_3_, or neuronal populations in the vicinity of the hotspot. Thus, our findings suggest that axonal activation occurred preferentially with the E-field aligned parallel to neuronal bundles at bending terminals, supported by earlier simulation studies [19]. This is in line with the observation that an increase in stimulation intensity seems to reduce the neuronal selectivity of the TMS pulse.

We observed that the MEP latency at 180° was marginally longer (0.7 ms) than at 0° with 110% of MT_0°_. Similar latency differences between 0° (posterior–anterior) and 180° (anterior–posterior) have been reported also with half-sine [53] and near-rectangular [54] E-field waveforms, like the one used in this study, different from the 2–3 ms longer latency for anterior–posterior than for posterior–anterior directions reported with conventional monophasic pulses [53]. During the brief falling part of our current waveform, the E-field has a relatively strong amplitude but reversed polarity compared to the E-field during the rising part of the current waveform. This relatively strong reversed E-field might also cause different neuronal effects, e.g., at the level of sodium channels and membrane time constants, than a conventional monophasic TMS pulse with a longer-lasting but weaker E-field during the current decay, as demonstrated with simulations of a biphasic electric stimulus [55]. Therefore, stimuli at 0° and 180° orientations with symmetric biphasic E-fields possibly excite similar direction-specific neuronal inputs with distinct thresholds, as recently described by Sommer et al. [54]. This mechanism likely explains the significantly different amplitude but similar latency between both orientations.

We should note that the relation between MEP parameters and stimulus orientation and intensity might differ depending on several factors, such as the pulse waveform [33,53,56] and the muscle under study [2,57]. The neuronal depolarization with near-rectangular E-field waveforms [31] with short current fall times (like ours) may be more selective, at least for predominantly larger fibers, than those with relatively long current fall times used in conventional monophasic TMS as in Refs. [53,58,59].

The neurophysiological implications from our results should be carefully interpreted as the estimated E-field orientation is based on the spherical head geometry used to design and calibrate the mTMS transducer, not on a realistic morphology. However, due to the symmetric and smooth change in MEP amplitude relative to the stimulus orientation (**Fig. 4C**), it is unlikely that spurious changes in the E-field maximum [21,23] in the cortex would affect the recorded MEPs.

The 2-coil transducer provides fine electronic control of the peak E-field orientation in the spherical head model. In the real cortex, local anisotropy and the complex conductivity structure in the vicinity, however, make the E-field rotate at a somewhat different rate than the magnetic field. In this context, individual MRI-based E-field modeling would help estimate the realized orientation and intensity of the peak E-field on the cortical surface.

## Acknowledgments

The authors would like to thank Ana Soto de la Cruz, Dr. Baran Aydogan, and Heikki Sinisalo for their assistance during the experiments, Dr. Tuomas Mutanen and Dr. Pantelis Lioumis for insightful discussions, and the Science-IT at Aalto University School of Science for the computational resources.

