## Supplementary Material for "TMS with fast and accurate electronic control: measuring the orientation sensitivity of corticomotor pathways"

#### Statistical analysis

The linear mixed-effects model residuals were inspected for critical deviations from normality with Q–Q plots, and homoscedasticity was inspected with a standard versus fitted values plot. Statistical analysis was performed in R 4.0 (R Core Team, Austria) using the *lme4* 1.1, *afex* 0.28 packages, and *emmeans* 1.5 packages.

### Supplementary figures

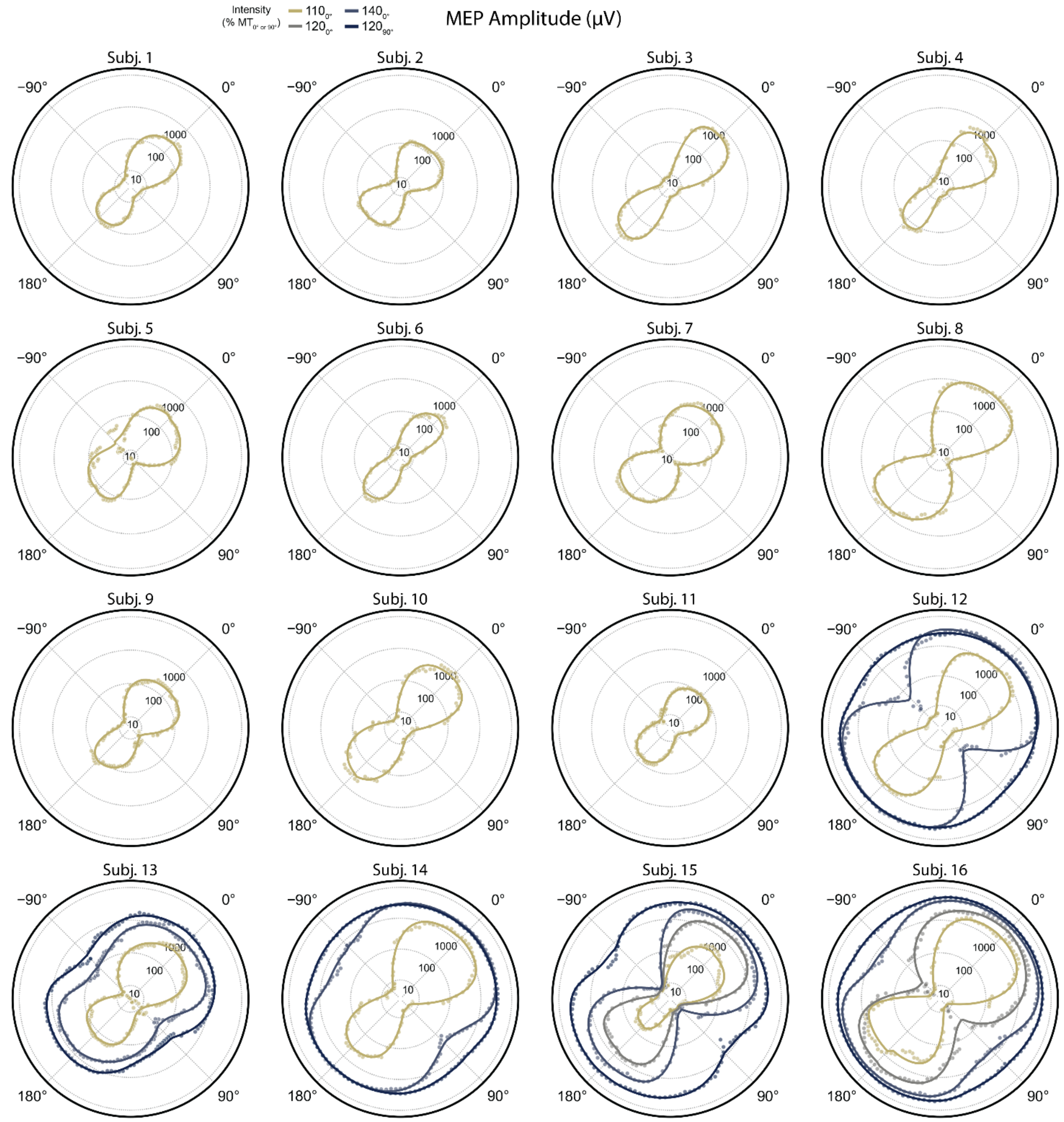


**Supplementary Figure S1: Motor evoked potential (MEP) amplitude as a function of the stimulus orientation and intensity for each subject.** The solid lines represent the logistic fit of the MEP amplitude (radial axis in a logarithmic scale) for each stimulation intensity in % of the resting motor threshold at 0° ($MT_{0°}$) and 90° ($MT_{90°}$). The dots represent the median MEP amplitude across the repeated trials for each stimulus condition.

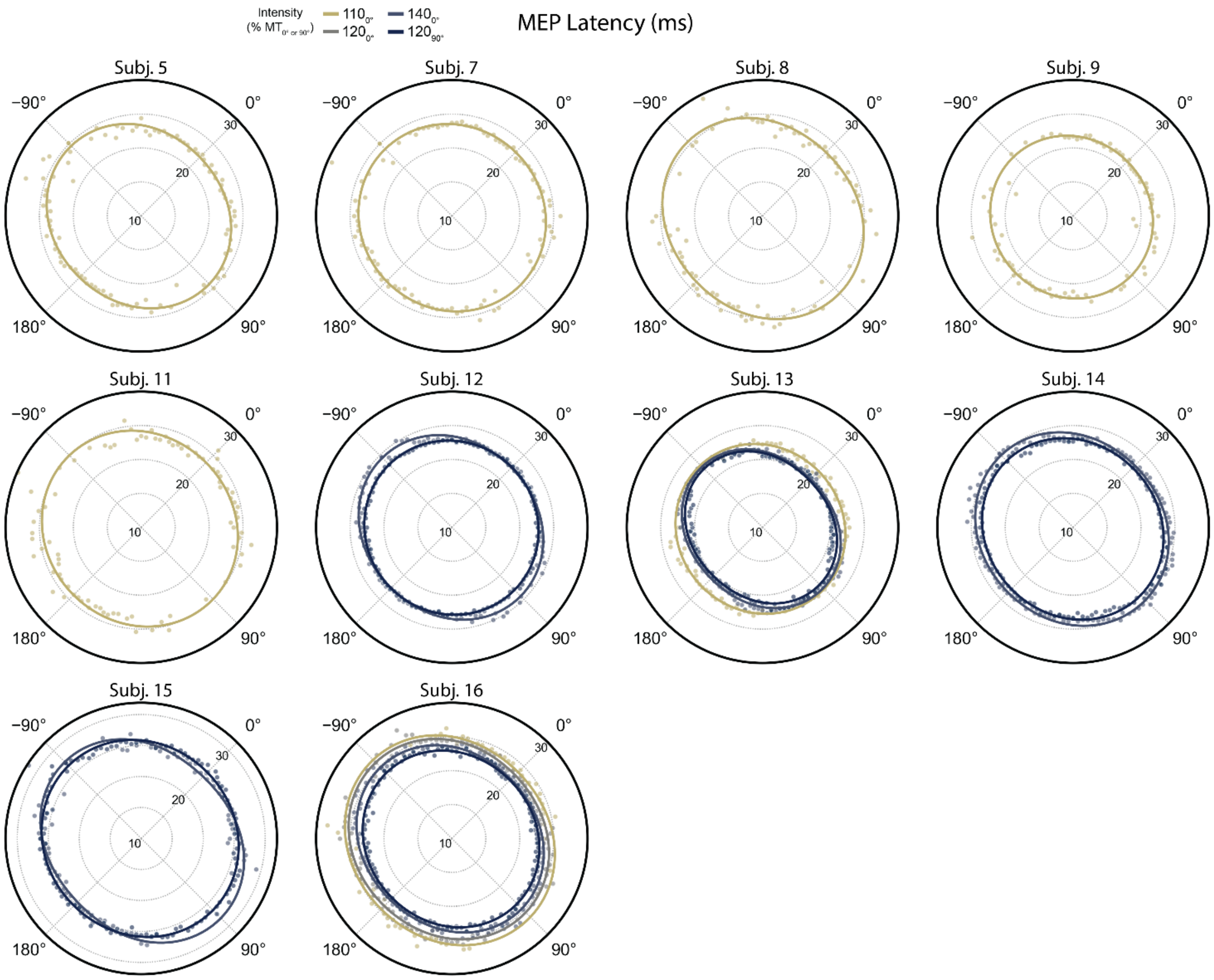


**Supplementary Figure S2: Motor evoked potential (MEP) latency as a function of the stimulus orientation and intensity for each subject.** The solid lines represent the trigonometric polynomial fit of the MEP latency for each stimulation intensity in % of the resting motor threshold at 0° ($MT_{0°}$) and 90° ($MT_{90°}$). The dots represent the median MEP amplitude across the repeated trials for each stimulus condition. We selected only subjects showing latencies across the entire range of orientations for each intensity (seven subjects in 110% $MT_{0°}$, one subject in 120% $MT_{0°}$, five subjects in 140% $MT_{0°}$ and 120% $MT_{90°}$).

### Supplementary tables

**Supplementary Table S1:** Type III Analysis of Variance Table with Satterthwaite's method for the mixed-effect model of the maximum motor evoked potential (MEP) amplitude. Fixed factors were the stimulation intensity (SI) and peak. Interaction between factors is represented as "×", and *p*-values in bold are smaller than the statistical significance level of 0.05.

| Effect | Degrees of freedom (numerator, denominator) | F-value | *p*-value |
|---|---|---|---|
| SI | (3, 34.4) | 56.5 | **< 0.001** |
| Peak | (1, 32.7) | 7.6 | **0.009** |
| SI × Peak | (3, 32.7) | 2.6 | 0.069 |

**Supplementary Table S2:** Multiple comparisons on maximum motor evoked potential (MEP) amplitude between stimulation intensities (SI). The results are presented as follows, for a fixed peak orientation, we compute the MEP amplitude ratio between the tested intensities. Each comparison has a standard error (SE), degrees of freedom (DoF), *t*-ratio, and *p*-value. The SI is given as a percentage of the orientation-specific resting motor threshold at 0° ($MT_{0°}$) and 90° ($MT_{90°}$). *p*-values in bold are smaller than the statistical significance level of 0.05.

| Peak | Comparison $SI_1$ / $SI_2$ | Amplitude ratio | SE | DoF | *t*-ratio | *p*-value |
|---|---|---|---|---|---|---|
| 0° | $110_{0°}$ / $120_{0°}$ | 0.27 | 0.09 | 34.22 | −3.98 | **0.001** |
| 0° | $110_{0°}$ / $140_{0°}$ | 0.22 | 0.05 | 34.94 | −6.65 | **< 0.001** |
| 0° | $110_{0°}$ / $120_{90°}$ | 0.18 | 0.04 | 34.94 | −7.65 | **< 0.001** |
| 0° | $120_{0°}$ / $140_{0°}$ | 0.81 | 0.28 | 33.27 | −0.61 | 0.625 |
| 0° | $120_{0°}$ / $120_{90°}$ | 0.65 | 0.22 | 33.27 | −1.27 | 0.296 |
| 0° | $140_{0°}$ / $120_{90°}$ | 0.80 | 0.20 | 33.03 | −0.90 | 0.458 |
| 180° | $110_{0°}$ / $120_{0°}$ | 0.29 | 0.09 | 34.22 | −3.80 | **0.001** |
| 180° | $110_{0°}$ / $140_{0°}$ | 0.13 | 0.03 | 34.94 | −9.15 | **< 0.001** |
| 180° | $110_{0°}$ / $120_{90°}$ | 0.09 | 0.02 | 34.94 | −10.53 | **< 0.001** |
| 180° | $120_{0°}$ / $140_{0°}$ | 0.44 | 0.15 | 33.27 | −2.43 | **0.037** |
| 180° | $120_{0°}$ / $120_{90°}$ | 0.32 | 0.11 | 33.27 | −3.34 | **0.004** |
| 180° | $140_{0°}$ / $120_{90°}$ | 0.73 | 0.18 | 33.03 | −1.25 | 0.296 |

**Supplementary Table S3:** Multiple comparisons on maximum motor evoked potential (MEP) amplitude between the peak orientations. The results are presented as follows, for a fixed stimulation intensity (SI), we compute the maximum MEP amplitude ratio between the peak orientations. Each comparison has a standard error (SE), degrees of freedom (DoF), *t*-ratio, and *p*-value. The SI is given as a percentage of the orientation-specific resting motor threshold at 0° ($MT_{0°}$) and 90° ($MT_{90°}$). *p*-values in bold are smaller than the statistical significance level of 0.05.

| SI % $MT_{0° or 90°}$ | Amplitude ratio Peak 0° / 180° | SE | DoF | *t*-ratio | *p*-value |
|---|---|---|---|---|---|
| $110_{0°}$ | 1.95 | 0.27 | 33.03 | 4.78 | **< 0.001** |
| $120_{0°}$ | 2.07 | 0.82 | 33.03 | 1.84 | 0.120 |
| $140_{0°}$ | 1.11 | 0.28 | 33.03 | 0.41 | 0.726 |
| $120_{90°}$ | 1.02 | 0.26 | 33.03 | 0.07 | 0.942 |

**Supplementary Table S4:** Type III Analysis of Variance Table with Satterthwaite's method for the mixed-effect model of the baseline motor evoked potential (MEP) amplitude. Fixed factors were the stimulation intensity (SI) and peak. Interaction between factors is represented as "×", and *p*-values in bold are smaller than the statistical significance level of 0.05.

| Effect | Degrees of freedom (numerator, denominator) | F-value | *p*-value |
|---|---|---|---|
| SI | (3, 45.1) | 78.6 | **< 0.001** |
| Peak | (1, 40.8) | 4.0 | 0.052 |
| SI × Peak | (3, 40.8) | 1.7 | 0.190 |

**Supplementary Table S5:** Multiple comparisons on baseline motor evoked potential (MEP) amplitude between stimulation intensities (SI). The results are presented as follows, for a fixed peak orientation, we compute the baseline MEP amplitude ratio between the tested intensities. Each comparison has a standard error (SE), degrees of freedom (DoF), *t*-ratio, and *p*-value. The SI is given as a percentage of the orientation-specific resting motor threshold at 0° ($MT_{0°}$) and 90° ($MT_{90°}$). *p*-values in bold are smaller than the statistical significance level of 0.05.

| Peak | Comparison $SI_1$ / $SI_2$ | Amplitude ratio | SE | DoF | *t*-ratio | *p*-value |
|---|---|---|---|---|---|---|
| 0° | $110_{0°}$ / $120_{0°}$ | 4.27 | 2.99 | 42.94 | 2.08 | 0.058 |
| 0° | $110_{0°}$ / $140_{0°}$ | 0.09 | 0.04 | 43.42 | −5.06 | **< 0.001** |
| 0° | $110_{0°}$ / $120_{90°}$ | 0.01 | < 0.01 | 43.42 | −10.13 | **< 0.001** |
| 0° | $120_{0°}$ / $140_{0°}$ | 0.02 | 0.02 | 38.47 | −5.07 | **< 0.001** |
| 0° | $120_{0°}$ / $120_{90°}$ | < 0.01 | < 0.01 | 38.47 | −8.22 | **< 0.001** |
| 0° | $140_{0°}$ / $120_{90°}$ | 0.09 | 0.05 | 34.97 | −4.30 | **< 0.001** |
| 180° | $110_{0°}$ / $120_{0°}$ | 0.56 | 0.39 | 42.94 | −0.84 | 0.502 |
| 180° | $110_{0°}$ / $140_{0°}$ | 0.10 | 0.05 | 43.42 | −4.93 | **< 0.001** |
| 180° | $110_{0°}$ / $120_{90°}$ | 0.01 | < 0.01 | 43.42 | −10.51 | **< 0.001** |
| 180° | $120_{0°}$ / $140_{0°}$ | 0.17 | 0.13 | 38.47 | −2.30 | **0.039** |
| 180° | $120_{0°}$ / $120_{90°}$ | 0.01 | 0.01 | 38.47 | −5.77 | **< 0.001** |
| 180° | $140_{0°}$ / $120_{90°}$ | 0.07 | 0.04 | 34.97 | −4.73 | **< 0.001** |

**Supplementary Table S6:** Multiple comparisons on motor evoked potential (MEP) baseline amplitude between the peak orientations. The results are presented as follows, for a fixed stimulation intensity (SI), we compute the baseline MEP amplitude ratio between the peak orientations. Each comparison has a standard error (SE), degrees of freedom (DoF), *t*-ratio, and *p*-value. The SI is given as a percentage of the orientation-specific resting motor threshold at 0° ($MT_{0°}$) and 90° ($MT_{90°}$). *p*-values in bold are smaller than the statistical significance level of 0.05.

| SI % $MT_{0° or 90°}$ | Amplitude ratio Peak 0° / 180° | SE | DoF | *t*-ratio | *p*-value |
|---|---|---|---|---|---|
| $110_{0°}$ | 0.93 | 0.29 | 34.97 | −0.23 | 0.871 |
| $120_{0°}$ | 0.12 | 0.11 | 34.97 | −2.4 | **0.035** |
| $140_{0°}$ | 0.99 | 0.55 | 34.97 | −0.02 | 0.983 |
| $120_{90°}$ | 0.78 | 0.43 | 34.97 | −0.45 | 0.751 |

**Supplementary Table S7:** Type III Analysis of Variance Table with Satterthwaite's method for the mixed-effect model of the centre orientation. Fixed factors were the stimulation intensity (SI) and peak. Interaction between factors is represented as "×", and *p*-values in bold are smaller than the statistical significance level of 0.05.

| **Effect** | **Degrees of freedom (numerator, denominator)** | **F-value** | ***p*-value** |
|---|---|---|---|
| SI | (3, 39.9) | 20.4 | **< 0.001** |
| Peak | (1, 31.0) | 14.8 | **< 0.001** |
| SI × Peak | (3, 31.0) | 0.9 | 0.432 |

**Supplementary Table S8:** Multiple comparisons on centre orientation between stimulation intensities (SI). The results are presented as follows, for a peak orientation, we compute the centre orientation difference between the tested intensities. Each comparison has a standard error (SE), degrees of freedom (DoF), *t*-ratio, and *p*-value. The SI is given as a percentage of the orientation-specific resting motor threshold at 0° ($MT_{0°}$) and 90° ($MT_{90°}$). *p*-values in bold are smaller than the statistical significance level of 0.05.

| **Peak** | **Comparison $SI_1 - SI_2$** | **Centre difference** | **SE** | **DoF** | ***t*-ratio** | ***p*-value** |
|---|---|---|---|---|---|---|
| 0° | $110_{0°} - 120_{0°}$ | 21.38 | 6.61 | 42.24 | 3.23 | **0.008** |
| 0° | $110_{0°} - 140_{0°}$ | 18.29 | 4.48 | 43.04 | 4.09 | **0.001** |
| 0° | $110_{0°} - 120_{90°}$ | 22.98 | 4.48 | 43.04 | 5.14 | **< 0.001** |
| 0° | $120_{0°} - 140_{0°}$ | −3.09 | 7.16 | 37.84 | −0.43 | 0.713 |
| 0° | $120_{0°} - 120_{90°}$ | 1.60 | 7.16 | 37.84 | 0.22 | 0.824 |
| 0° | $140_{0°} - 120_{90°}$ | 4.69 | 5.25 | 34.71 | 0.89 | 0.465 |
| 180° | $110_{0°} - 120_{0°}$ | 6.81 | 6.61 | 42.24 | 1.03 | 0.412 |
| 180° | $110_{0°} - 140_{0°}$ | 18.31 | 4.48 | 43.04 | 4.09 | **0.001** |
| 180° | $110_{0°} - 120_{90°}$ | 21.13 | 4.48 | 43.04 | 4.72 | **< 0.001** |
| 180° | $120_{0°} - 140_{0°}$ | 11.50 | 7.16 | 37.84 | 1.61 | 0.187 |
| 180° | $120_{0°} - 120_{90°}$ | 14.32 | 7.16 | 37.84 | 2.00 | 0.105 |
| 180° | $140_{0°} - 120_{90°}$ | 2.82 | 5.25 | 34.71 | 0.54 | 0.679 |

**Supplementary Table S9:** Multiple comparisons on centre orientation between the peak orientations. The results are presented as follows, for a fixed stimulation intensity (SI), we compute the centre difference between the peak orientations. Each comparison has a standard error (SE), degrees of freedom (DoF), *t*-ratio, and *p*-value. The SI is given as a percentage of the orientation-specific resting motor threshold at 0° ($MT_{0°}$) and 90° ($MT_{90°}$). *p*-values in bold are smaller than the statistical significance level of 0.05.

| **SI % $MT_{0° or 90°}$** | **Centre difference Peak 0° − 180°** | **SE** | **DoF** | ***t*-ratio** | ***p*-value** |
|---|---|---|---|---|---|
| $110_{0°}$ | −6.97 | 2.93 | 34.71 | −2.38 | 0.053 |
| $120_{0°}$ | −21.54 | 8.3 | 34.71 | −2.6 | **0.037** |
| $140_{0°}$ | −6.96 | 5.25 | 34.71 | −1.33 | 0.282 |
| $120_{90°}$ | −8.83 | 5.25 | 34.71 | −1.68 | 0.181 |

**Supplementary Table S10:** Type III Analysis of Variance Table with Satterthwaite's method for the mixed-effect model of the sigma. Fixed factors were the stimulation intensity (SI) and peak. Interaction between factors is represented as "×", and *p*-values in bold are smaller than the statistical significance level of 0.05.

| Effect | Degrees of freedom (numerator, denominator) | F-value | *p*-value |
|---|---|---|---|
| SI | (3, 38.5) | 0.44 | 0.727 |
| Peak | (1, 30.4) | 12.0 | **0.001** |
| SI × Peak | (3, 30.4) | 2.7 | 0.063 |

**Supplementary Table S11:** Multiple comparisons on sigma between stimulation intensities (SI). The results are presented as follows, for a peak orientation, we compute the sigma difference between the tested intensities. Each comparison has a standard error (SE), degrees of freedom (DoF), *t*-ratio, and *p*-value. The SI is given as a percentage of the orientation-specific resting motor threshold at 0° ($MT_{0°}$) and 90° ($MT_{90°}$). *p*-values in bold are smaller than the statistical significance level of 0.05.

| Peak | Comparison $SI_1 - SI_2$ | Sigma difference | SE | DoF | *t*-ratio | *p*-value |
|---|---|---|---|---|---|---|
| 0° | $110_{0°} - 120_{0°}$ | 0.04 | 0.02 | 39.97 | 1.82 | 0.255 |
| 0° | $110_{0°} - 140_{0°}$ | 0.00 | 0.02 | 41.55 | −0.02 | 0.980 |
| 0° | $110_{0°} - 120_{90°}$ | 0.03 | 0.02 | 41.55 | 2.06 | 0.243 |
| 0° | $120_{0°} - 140_{0°}$ | −0.04 | 0.03 | 36.07 | −1.71 | 0.255 |
| 0° | $120_{0°} - 120_{90°}$ | −0.01 | 0.03 | 36.07 | −0.39 | 0.889 |
| 0° | $140_{0°} - 120_{90°}$ | 0.03 | 0.02 | 34.00 | 1.81 | 0.255 |
| 180° | $110_{0°} - 120_{0°}$ | −0.01 | 0.02 | 39.97 | −0.35 | 0.889 |
| 180° | $110_{0°} - 140_{0°}$ | 0.01 | 0.02 | 41.55 | 0.52 | 0.879 |
| 180° | $110_{0°} - 120_{90°}$ | −0.01 | 0.02 | 41.55 | −0.92 | 0.642 |
| 180° | $120_{0°} - 140_{0°}$ | 0.02 | 0.03 | 36.07 | 0.66 | 0.824 |
| 180° | $120_{0°} - 120_{90°}$ | −0.01 | 0.03 | 36.07 | −0.26 | 0.889 |
| 180° | $140_{0°} - 120_{90°}$ | −0.02 | 0.02 | 34.00 | −1.25 | 0.458 |

**Supplementary Table S12:** Multiple comparisons on sigma between the peak orientations. The results are presented as follows, for a fixed stimulation intensity (SI), we compute the sigma difference between the peak orientations. Each comparison has a standard error (SE), degrees of freedom (DoF), *t*-ratio, and *p*-value. The SI is given as a percentage of the orientation-specific resting motor threshold at 0° ($MT_{0°}$) and 90° ($MT_{90°}$). *p*-values in bold are smaller than the statistical significance level of 0.05.

| SI % $MT_{0° or 90°}$ | Sigma difference Peak 0° − 180° | SE | DoF | *t*-ratio | *p*-value |
|---|---|---|---|---|---|
| $110_{0°}$ | −0.01 | 0.01 | 34 | −1.23 | 0.458 |
| $120_{0°}$ | −0.06 | 0.03 | 34 | −2.18 | 0.243 |
| $140_{0°}$ | 0.00 | 0.02 | 34 | −0.21 | 0.889 |
| $120_{90°}$ | −0.06 | 0.02 | 34 | −3.27 | **0.039** |

**Supplementary Table S13:** Type III Analysis of Variance Table with Satterthwaite's method for the mixed-effect model of the motor evoked potential latency. Fixed factors were the stimulation intensity (SI) and peak. Interaction between factors is represented as "×", and *p*-values in bold are smaller than the statistical significance level of 0.05.

| Effect | Degrees of freedom (numerator, denominator) | F-value | *p*-value |
|---|---|---|---|
| SI | (2, 48.1) | 49.0 | **< 0.001** |
| Peak | (3, 46.9) | 93.6 | **< 0.001** |
| SI × Peak | (6, 46.9) | 3.5 | 0.006 |

**Supplementary Table S14:** Multiple comparisons on motor evoked amplitude (MEP) latency between stimulation intensities (SI). The results are presented as follows, for a fixed peak orientation, we compute the MEP latency difference between the tested intensities. Each comparison has a standard error (SE), degrees of freedom (DoF), *t*-ratio, and *p*-value. The SI is given as a percentage of the orientation-specific resting motor threshold at 0° ($MT_{0°}$) and 90° ($MT_{90°}$). *p*-values in bold are smaller than the statistical significance level of 0.05.

| Peak | Comparison $SI_1$−$SI_2$ | Latency difference | SE | DoF | *t*-ratio | *p*-value |
|---|---|---|---|---|---|---|
| 0° | $110_{0°} - 140_{0°}$ | 1.71 | 0.29 | 47.99 | 6 | **< 0.001** |
| 0° | $110_{0°} - 120_{90°}$ | 1.67 | 0.29 | 47.99 | 5.84 | **< 0.001** |
| 0° | $140_{0°} - 120_{90°}$ | −0.05 | 0.26 | 47.00 | −0.18 | 0.857 |
| 90° | $110_{0°} - 140_{0°}$ | 0.83 | 0.29 | 47.99 | 2.9 | **0.008** |
| 90° | $110_{0°} - 120_{90°}$ | 2.00 | 0.29 | 47.99 | 7.00 | **< 0.001** |
| 90° | $140_{0°} - 120_{90°}$ | 1.17 | 0.26 | 47.00 | 4.43 | **< 0.001** |
| 180° | $110_{0°} - 140_{0°}$ | 1.84 | 0.29 | 47.99 | 6.44 | **< 0.001** |
| 180° | $110_{0°} - 120_{90°}$ | 1.99 | 0.29 | 47.99 | 6.96 | **< 0.001** |
| 180° | $140_{0°} - 120_{90°}$ | 0.15 | 0.26 | 47.00 | 0.56 | 0.668 |
| −90° | $110_{0°} - 140_{0°}$ | 0.84 | 0.29 | 47.99 | 2.93 | **0.007** |
| −90° | $110_{0°} - 120_{90°}$ | 1.85 | 0.29 | 47.99 | 6.47 | **< 0.001** |
| −90° | $140_{0°} - 120_{90°}$ | 1.01 | 0.26 | 47.00 | 3.82 | **0.001** |

**Supplementary Table S15:** Multiple comparisons on motor evoked potential (MEP) latency between the peak orientations. The results are presented as follows, for a fixed stimulation intensity (SI), we compute the sigma difference between the peak orientations. Each comparison has a standard error (SE), degrees of freedom (DoF), *t*-ratio, and *p*-value. The SI is given as a percentage of the orientation-specific resting motor threshold at 0° ($MT_{0°}$) and 90° ($MT_{90°}$). *p*-values in bold are smaller than the statistical significance level of 0.05.

| **SI % $MT_{0° or 90°}$** | **Comparison $Peak_1 - Peak_2$** | **Latency difference** | **SE** | **DoF** | ***t*-ratio** | ***p*-value** |
|---|---|---|---|---|---|---|
| $110_{0°}$ | 0° − 90° | −1.77 | 0.22 | 47 | −7.91 | **< 0.001** |
| $110_{0°}$ | 0° − 180° | −0.64 | 0.22 | 47 | −2.85 | **0.008** |
| $110_{0°}$ | 0° − (−90°) | −1.67 | 0.22 | 47 | −7.50 | **< 0.001** |
| $110_{0°}$ | 90° – 180° | 1.13 | 0.22 | 47 | 5.06 | **< 0.001** |
| $110_{0°}$ | 90° – (−90°) | 0.09 | 0.22 | 47 | 0.41 | 0.753 |
| $110_{0°}$ | 180° − (−90°) | −1.04 | 0.22 | 47 | −4.64 | **< 0.001** |
| $140_{0°}$ | 0° − 90° | −2.65 | 0.26 | 47 | −10.04 | **< 0.001** |
| $140_{0°}$ | 0° − 180° | −0.51 | 0.26 | 47 | −1.94 | 0.072 |
| $140_{0°}$ | 0° − (−90°) | −2.55 | 0.26 | 47 | −9.65 | **< 0.001** |
| $140_{0°}$ | 90° – 180° | 2.14 | 0.26 | 47 | 8.10 | **< 0.001** |
| $140_{0°}$ | 90° – (−90°) | 0.10 | 0.26 | 47 | 0.38 | 0.753 |
| $140_{0°}$ | 180° − (−90°) | −2.04 | 0.26 | 47 | −7.71 | **< 0.001** |
| $120_{90°}$ | 0° − 90° | −1.43 | 0.26 | 47 | −5.43 | **< 0.001** |
| $120_{90°}$ | 0° − 180° | −0.32 | 0.26 | 47 | −1.20 | 0.282 |
| $120_{90°}$ | 0° − (−90°) | −1.49 | 0.26 | 47 | −5.65 | **< 0.001** |
| $120_{90°}$ | 90° – 180° | 1.12 | 0.26 | 47 | 4.22 | **< 0.001** |
| $120_{90°}$ | 90° – (−90°) | −0.06 | 0.26 | 47 | −0.22 | 0.852 |
| $120_{90°}$ | 180° − (−90°) | −1.18 | 0.26 | 47 | −4.45 | **< 0.001** |
